# Interactions Between Native Soil Microbiome and a Synthetic Microbial Community Reveals Bacteria with Persistent Traits

**DOI:** 10.1101/2025.04.08.647796

**Authors:** Jessica M. Velte, Sameerika Dissanayaka Mudiyanselage, Olivia F. Hofmann, Sonny T.M. Lee, Jose Huguet-Tapia, Mariza Miranda, Samuel J. Martins

## Abstract

Synthetic microbial communities (SynComs) are curated microbial groups designed to enhance plant growth or disease resistance by augmenting soil microbiomes. Attaining SynCom stability in the presence of native soil communities remains a key challenge. This study investigated the survival, persistence, and chemical interactions of a SynCom with a native soil microbial community using a transwell system that spatially constrains bacteria while permitting chemical interactions. The SynCom, composed of six compatible Pseudomonas species identified through whole-genome sequencing, was analyzed for antagonistic interactions with native microbes over time and assessed using biomass and viability measurements. Over time, the SynCom exhibited reduced growth in the presence of native soil microbes compared to the SynCom not exposed to the native microbes. Flow cytometry analysis showed an 81% reduction of live cells for the persistent strain in the presence of native microbes and a 78% and 99% increase in dead and unstained cells, respectively. Compared to a non-persistent strain, one persistent SynCom strain showed lower metabolic utilization across five key compound classes: polymers, carboxylic acids, amino acids, amines, and phenols when exposed to the native soil microbes. These findings underscore the importance of understanding complex SynCom-environment interactions to enhance SynCom stability and optimize *in situ* applications.

**Importance:** Synthetic Microbial Communities, or SynComs, are an emerging technology that can potentially augment plant health. Still, their application *in situ* depends on deciphering the complex interactions between SynCom microbes and native microbial communities. This study provides insight into several *Pseudomonas* strains displaying persistent characteristics, which makes these bacteria promising candidates for SynCom stability in implanted environments. Understanding the persistent traits of these bacteria is a vital advancement in SynCom technology, and an important next step toward implementing SynComs in agricultural systems.

## Introduction

Synthetic microbial communities (SynComs) are curated groups of microbes designed to enhance specific microbial functions. SynComs can be used to augment the plant soil microbiome and improve plant growth or disease resistance. For instance, Mažylytė et al. (2024) demonstrated that SynComs comprising beneficial rhizobacteria, including *Bacillus*, *Pseudomonas*, and *Streptomyces*, promoted nutrient uptake in *Triticum aestivum* by enhancing root growth and improving nutrient availability. Similarly, Yang et al. (2021) found that a bacterial consortium consisting of *Stenotrophomonas rhizophila*, *Xanthomonas retroflexus*, *Microbacterium oxydans*, and *Paenibacillus amylolyticus* induced drought tolerance in *Arabidopsis thaliana*. This effect was attributed to sustained chlorophyll content and the activation of endogenous abscisic acid (ABA) signaling pathways, both critical for plant adaptation under water stress. SynComs comprising different bacterial genera have been specially designed for a variety of reasons, such as to enhance tomato growth and suppress symptoms of *Fusarium* wilt (Tsolakidou et al. 2019) or suppress *Rhizoctonia solani* infections on wheat (Yin et al. 2022).

However, among the challenges regarding utilizing SynComs is their lack of persistence in the environment where they are applied (Marín et al. 2022; Ahmad et al. 2024). To ensure longevity and retain both function and benefit in applied environments, the microorganisms comprising the SynCom must not only be compatible (Martins et al. 2023) but also successfully integrate with the native soil microbiome.

*Pseudomonas* spp., a prominent group of Gram-negative bacteria within plant-growth promoting rhizobacteria (PGPR), exhibit strong antimicrobial and biocontrol properties (Masschelein et al. 2017; Wang et al. 2021). Commonly found in the soil in a narrow area around the roots, known as the rhizosphere, *Pseudomonas* spp. produce various secondary metabolites which inhibit pathogens, including fungi, oomycetes, bacteria, viruses, and protozoans (Masschelein et al. 2017; Zhao et al. 2019). In a previous study aiming to study the effects of a dysbiotic or imbalanced microbial community, induced by antibiotic application of streptomycin in the citrus rhizosphere, it was found that *Pseudomonas* decreased and photosynthesis reduced, making *Pseudomonas* a valuable addition (Ketehouli et al. 2024). These characteristics make *Pseudomonas* spp. valuable candidates for inclusion in SynComs. While promising in strategy, SynComs face challenges in implementation. Notably, the persistence of SynCom members within the environment where they are applied is a critical factor to be considered while designing a SynCom with community resilience (Marín et al. 2022; Jiang et al. 2022; Ahmad et al. 2024; Martins et al. 2023). To survive and establish in the environment, the SynCom members need to be compatible among themselves (Martins et al. 2023) and successfully merge with the native soil microbiome. Microbial interactions mediated by allelochemicals play a significant role in these dynamics, as certain microbes can enter a state of reduced metabolic activity and growth, known as persistence, to survive unfavorable conditions (Fonseca-García et al. 2024; Lewis, 2007). Studying the microbial interactions among both SynCom members and the native soil microbial communities is an additional challenge in successfully implementing SynComs. Addressing how donor and recipient microbes respond to each other’s presence via metabolic interactions is key to identifying successful pairings of synthetic microbial communities with host microbiomes.

In this study, we hypothesized that synthetic microbial communities are reduced under perturbation in the presence of natural community vs. in the absence of natural community and assessed via transwell system, and some microbes are more affected than others.

The objectives of this study were to: (1) assess how a synthetic microbial community (SynCom) interacts with a native soil microbiome over time using a transwell system.; (2) evaluate possible persistence mechanisms of representative SynCom members assessed viability via flowcytometry and metabolic activity; (3) characterize the microbial composition and functional pathways of the native soil microbiome via shotgun metagenomic analysis, providing insight into SynCom interactions with native soil microbiome.

## Material and Methods

### Bacterial Strain collection and compatibility assessment

Bacterial strains (n=6) of the genus *Pseudomonas* isolated from citrus rhizosphere located in Citra, FL (Elevation: 2 m, 29**°** 24’38”, N 82**°** 8’57” W), were plated on King’s B (KB) media and tested for their compatibility among themselves using 10% Tryptic Soy Agar (TSA). For the compatibility test, single colonies from each strain were grown in liquid King’s B medium for 24 h at 28**°**C, and a single colony was suspended in 1 mL of sterile deionized water to standardize inoculum density. For each bacterial strain, 100 μL of the suspension was pipetted onto a plate with 10% TSA media and spread to make a bacterial lawn. The suspension was dried for approximately 10 mins, and then 10 μL of each bacterial suspension was pipetted onto the plate in discrete, untouching spots. Plates were incubated at 28°C for 24 h, and the compatibility of the bacteria was determined by observing zones of inhibition.

### Genotypic characterization via whole genome sequencing (WGS)

Genomic DNA was extracted from the six bacterial strains using the Quick-DNA Fecal/Soil Microbe MiniPrep Kit (ZYMO). DNA quality and quantity were checked using a Nanodrop spectrophotometer and sent to SeqCenter (Pittsburgh, PA) for sequencing. Illumina sequencing libraries were prepared using an Illumina DNA Prep kit with custom 10 bp unique dual indices. Sequencing was performed on an Illumina NovaSeq X Plus, generating 2×151 bp paired-end reads and processed using bcl-convert for demultiplexing, quality control, and adapter trimming. Assemblies were performed with Unicycler (v0.5.0), and assembly statistics were generated using QUAST (Mikheenko et al. 2018). Annotation was completed using Bakta (v1.8.1).

### Transwell System

The extraction of the soil microbiome as a soil slurry followed the techniques from Ketehouli et al. (2024) and Zhou et al. (2019) with some modifications. Three citrus plants in Citra, FL (19 m elevation, 29**°** 24’38” N, 82**°** 8’57” W) were used to collect the rhizosphere samples from roots in the top 10-20 cm of soil (Jiang et al. 2022) per plant. Soil collected from the site was sent to the UF/IFAS Soil Testing Services for soil chemical analysis and is presented in Table 1.

**Table 1.**
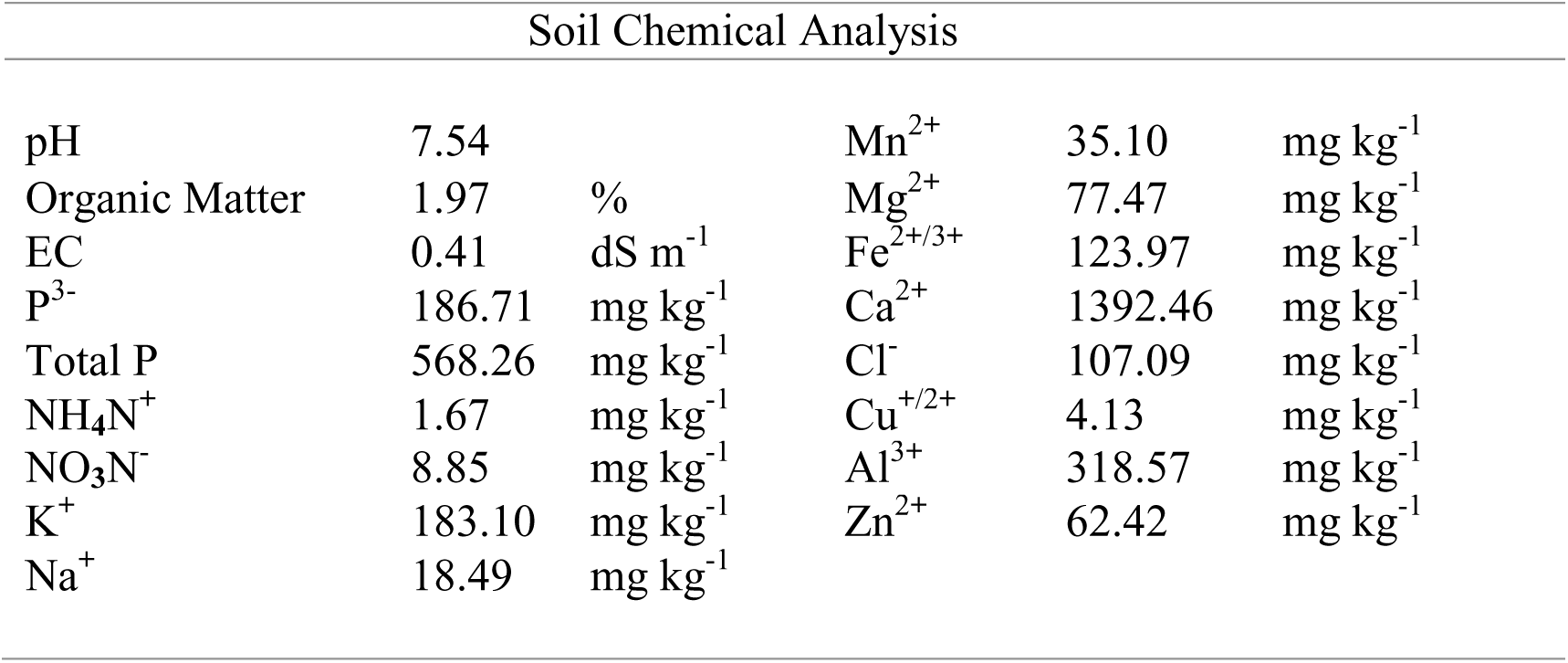
Assessment of soil chemical parameters, including macro- and micronutrients, pH, organic matter, and electrical conductivity (EC) of the soil used for the native soil microbe slurry.

The roots in the collected samples were shaken to remove extraneous soil in the root zone. The remaining soil on roots was collected by removing soil from the root surface through vigorous shaking and washing in [1mM] phosphate buffered saline (PBS). Fifteen grams of rhizosphere soil was mixed in 150 mL of PBS, excluding roots, in a 1:10 ratio (Mendes et al.2011). The mixture was blended three times for 60 s for homogenization to be representative of the native soil microbial community (Jiang et al. 2022). The soil suspension was then passed through a 200-mesh (0.07 mm) screen sieve to remove any remaining large particulate matter. At the beginning of each trial, three tubes of 1 mL of the soil slurry were collected for shotgun metagenomics and stored at −80°C until further processing.

To prepare the SynCom experiment, the aforementioned six bacterial isolates were grown on KB media. Single colonies of each bacteria were grown in liquid KB medium and incubated at 50 rpm, 25**°**C for 24h. Pre-addition to the plate, individual numbers of the SynCom were diluted in M9 minimal salts medium and OD_600_ were adjusted to have between 0.015 ±0.005 using a spectrophotometer (Bio-Rad, SmartSpec 3000 UV/Vis Spectrophotometer, Bio-Rad Laboratories, Hercules, CA, United States) were used to set up the SynCom experiment.

To assemble the transwell system, 96 well-plates with 0.22 µm filtered bottoms (Millipore®, Cat. No. MAGVS2210) were connected to a shared basal reservoir (Corning, Cat. No. 3383) to examine the interactions between the SynCom composed of 6 strains of *Pseudomonas* and native soil microbial community consisting of the soil slurry according to the Chodkowski and Shade (2017, 2024) set up. The filter of well H12 was removed using a sterile razor blade and forceps. The unfiltered well was used to carefully dispense 40 mL of M9 minimal salts medium or homogenized soil slurry in each plate, according to the treatment type. 200 µL of each bacterial suspension comprising the synthetic microbial community were pipetted into respective wells.

Placement of each bacterial suspension was based on a randomized selection of the wells using a custom R script created by Chodkowski and Shade (2017) and adapted to suit the number of bacteria used to create the synthetic microbial community in this study (RandomArray.R [see the GitHub repository]). Two treatments were used: 1) plate containing six bacteria in a randomized arrangement in the well plate (SynCom) and with the soil slurry at the bottom reservoir; 2) plate containing the same SynCom arrangement, however with the reservoir filled with M9 minimal medium as a control. Fifteen wells of each of the six bacteria were used alongside five blanks composed of M9 medium. The experiment was repeated, and each trial had a different randomized placement of the SynCom bacteria. To prevent contamination, each well plate was sealed with a Bio-Rad Microseal ‘B’ Seal (Cat. No. MSB1001) and parafilm. The microseal and parafilm were replaced at each time point. Optical density and plated viability data of the interacting microbes in the system was collected at 12.5 h, 25 h, 45 h, and 65 h based on growth stages of *Pseudomonas* spp.

### Bacterial growth in the transwell system

For each optical density reading for the previously defined time points, 10 µL of suspension from each well was pipetted into a sterile flat-bottomed 96 well-plate and mixed with 90 µL of PBS. OD was measured in a Bio-Rad iMark microplate reader (Cat. No. 1681130) at 595 nm. Pathway correction embedded in the reader software was applied to account for the 100 µL suspension volume in each well. To test bacterial viability, serial dilutions of the six bacterial strains ranged from 10^−5^ to 10^−7^, depending on the time point. A volume of 100 µL of the diluted bacterial suspension was spread evenly on plates with KB agar. Plates were incubated at 30**°**C for 24 h before colony forming units (CFU) counting using OpenCFU (Geissmann, 2013). Colony forming unit counts (CFUs mL^-1^) were normalized using log transformations for comparison.

### SynCom metabolism assessment

The carbon metabolism of two *Pseudomonas* spp. (SMV71 and SMP6) that showed the least and greatest difference of growth compared to the control at the last timepoint measured (65 h) in the transwell system test were selected for this metabolism assay. The bacteria growth was measured before and after 95 h of exposure to the native soil microbes using the Biolog EcoPlate^TM^ (NC0410462). To prepare the EcoPlates, 15 µL of bacterial suspensions (OD = 0.015 ± 0.005) were combined with 135 µL M9 minimal salts medium. The bacterial suspension for the pre- and post-native microbe exposure experiments was obtained from an M9 liquid culture and the transwell plate. SMV71 and SMP6 were inoculated into 31 carbon-containing EcoPlate wells. The EcoPlates were incubated at 28°C, 50 rpm for three days, and carbon utilization of each *Pseudomonas* spp. was measured at evenly spaced intervals as described previously for the transwell system assay. All measurements were performed in triplicate across three EcoPlates following a randomized block design.

### Flow cytometry analysis

Bacterial viability for persistent traits using flow cytometer was performed at the Cytometry Core (RRID:SCR_019119) - ICBR/UF. Strain SMV71, which displayed persistent traits in viability analysis, were sampled from the SynCom bacteria at 65 h from each treatment. SMP6 was also selected. For each strain type, 10 µL of three wells containing the bacteria were combined with 270 µL of 0.2 µm filtered PBS for a total volume of 300 µL of representative sample for the first replicate. Two additional replicates were prepared from a randomized assortment of wells, using samples from a total of nine different wells per bacterial strain per treatment.

*Pseudomonas protegens* strain SMP6 from culture was used for live and dead control groups at the same the concentration and solvent of the tested samples from experimentation. 500 µL of the total suspension of live bacteria was boiled at 100 °C for 15 min to be used as a control for dead bacteria.

Bacteria were stained at an optical density of 0.05 to 0.1 using the LIVE/DEAD™ BacLight™ Bacterial Viability and Counting Kit (Thermo Fischer Scientific, Cat. No. L34856). Propidium iodide (PI) and SYTO-9 nucleic acid stains were titrated and used in a 1:1 ratio of 0.2 µL per 1 mL of sample. Samples were diluted 1:9 using 0.2 µm filtered PBS and analyzed on a Cytoflex LX (Beckman Coulter). The detection threshold setting was adjusted to 2000 to appropriately detect bacterial cells. The PI was read using the PE channel and the SYTO-9 using the FITC channel. Bleach and water were run between samples to avoid carryover. 10 µL of each sample was recorded.

### Shotgun metagenomic analysis of the soil slurry microbiome

At 64 h after the transwell experiment set up, three samples of 1 mL soil slurry were collected for comparative metagenomics testing to 12.5 h samples. Genomic DNA was extracted from the soil slurry samples using the Quick-DNA Fecal/Soil Microbe MiniPrep Kit (ZYMO, Cat. No. D6010) following the manufacturer’s instructions. DNA concentration was estimated with Qubit3.0 Fluorometer, and the quality of the DNA was examined with Biorad Universal Hood Gel Doc XR UV Transilluminator 200VA system. Extracted gDNA was sent for metagenomic sequencing at SeqCenter (Pittsburgh, PA). Illumina sequencing libraries were prepared using the tagmentation-based and PCR-based Illumina DNA Prep kit and custom 10 bp unique dual indices (UDI) from IDT, targeting an insert size of 280 bp. Sequencing was performed on an Illumina NovaSeq X Plus sequencer in multiplexed shared-flow-cell runs, generating 2×151 bp paired-end reads. Demultiplexing, quality control, and adapter trimming were conducted using bcl-convert1 (v4.2.4).

Quality control of the raw sequencing data was assessed using FastQC (v0.12.1) (Chen et al. 2018). Based on the FastQC results, reads were trimmed to remove low-quality bases (<20 bases), and to remove adapter sequences 15 bases from both the 5’ and 3’ ends of each read were also trimmed using Trim Galore (v0.6.10) (Krueger F. 2015).

Paired-end-reads were concatenated and analyzed using MetaPhlAn 4.0 (Blanco-Míguez et al. 2023) for taxonomic annotation and relative abundance estimation. Data visualization was performed using heatmaps generated in R (version 4.4.1).

Functional profiling was conducted with HUMAnN v3.0 (Beghini et al. 2021), and pathway abundance data were used for downstream analyses. MicrobiomeAnalyst (https://www.microbiomeanalyst.ca/) was utilized for principal component analysis (PCA) plots at the initial timepoint and at 65 h of the treatment for the soil slurry microbiome. A total of 6,631,094 read counts were filtered based on default parameters. Additionally, 445 low-abundance features were removed based on prevalence thresholds, and 685 low-variance features were eliminated using interquartile range filtering. The normalized dataset was used for clustering analyses, including principal component analysis (PCA).

### Statistics and visualization

Statistical analysis for the OD, viability, and flow cytometry readings was performed using SigmaPlot 15.0 (Systat Software Inc., San Jose, CA, USA) software. Data normality was assessed using the Shapiro-Wilk normality test. If normality failed, the Wilcoxon Signed-Rank Test was used to compare medians of paired groups. Where normality passed, a paired t-test was used to test the mean differences between the treatments. A significance threshold of p ≤ 0.05 was used for result interpretation EcoPlate data analysis was carried out according to Sun et al. (2024) with some modifications. To elaborate, each EcoPlate block contained a control well inoculated with bacteria in M9 medium without exposure to additional carbon sources. The OD_595_ values for all 31 carbon-containing wells were corrected by subtracting the OD_595_ value of the control well from the OD_595_ of each carbon-containing well at every timepoint. Following Asif et al. (2024), any negative values were assumed to be zero, and plate readings greater than 3.5, the instrument’s detection limit, were considered 3.5. Average well-color development was calculated for each subset of carbon compounds using the equation described in Asif et al. (2024). The resulting sequences of the whole genome were analyzed from annotated short-read assemblies. Whole genome sequences were processed to determine taxonomic identification using the Type Strain Genome Server (TYGS) webserver to analyze digital DNA-DNA hybridization (dDDH) and pyani for average nucleotide identity (ANI). The ANI was performed using pyani version 0.2.12 (Pritchard et al. 2016). The chosen output for analysis was ANIm, which utilizes MUMer for pairwise genomic comparison. Whole genome sequences were further analyzed for prediction of secondary metabolites using biosynthetic gene clusters via antiSMASH version 7.1.0 (Medema et al. 2011).

The sequencing data supporting this study have been deposited in the NCBI database under Bioproject ID PRJNA1204327. Biosample accessions are SAMN46018761– SAMN46018766. Genome assemblies can be accessed via GenBank under the following accession numbers: JBKGGT000000000, JBKGGU000000000, JBKGGV000000000,JBKGGW000000000, JBKGGX000000000, and JBKGGY000000000. The associated organisms include *Pseudomonas* sp. SMV7, *Pseudomonas* sp. SMV71, *Pseudomonas* sp. SMSB3, *Pseudomona*s sp. SMN11, *Pseudomonas* sp. SMN5, and *Pseudomonas protegens* SMP6.

Metagenomics data is also available under the same Bioproject ID PRJNA1204327. Biosample accessions from SAMN46562606 to SAMN46562617.

## Results

### SynCom design and compatibility

Analysis of compatibility was based on visual observation for zones of inhibition on 10% TSA, and no visible zones of inhibition were observed between the bacteria strains. Bacterial colony growth did not appear to have abnormal color changes or morphological alterations and retained consistent size. Thus, the six strains were determined to be compatible.

### Molecular characterization and genomic comparison

Whole genome sequences were processed to determine taxonomic identification using the TYGS webserver. All results suggested that the strains were from the genus *Pseudomonas*. Confidence was analyzed through maximum likelihood tree construction and low *δ* value (Table 2). From the SynCom strains, strain SMP6 matched the species *P. protegens* (Fig. 1).

**Fig. 1.**
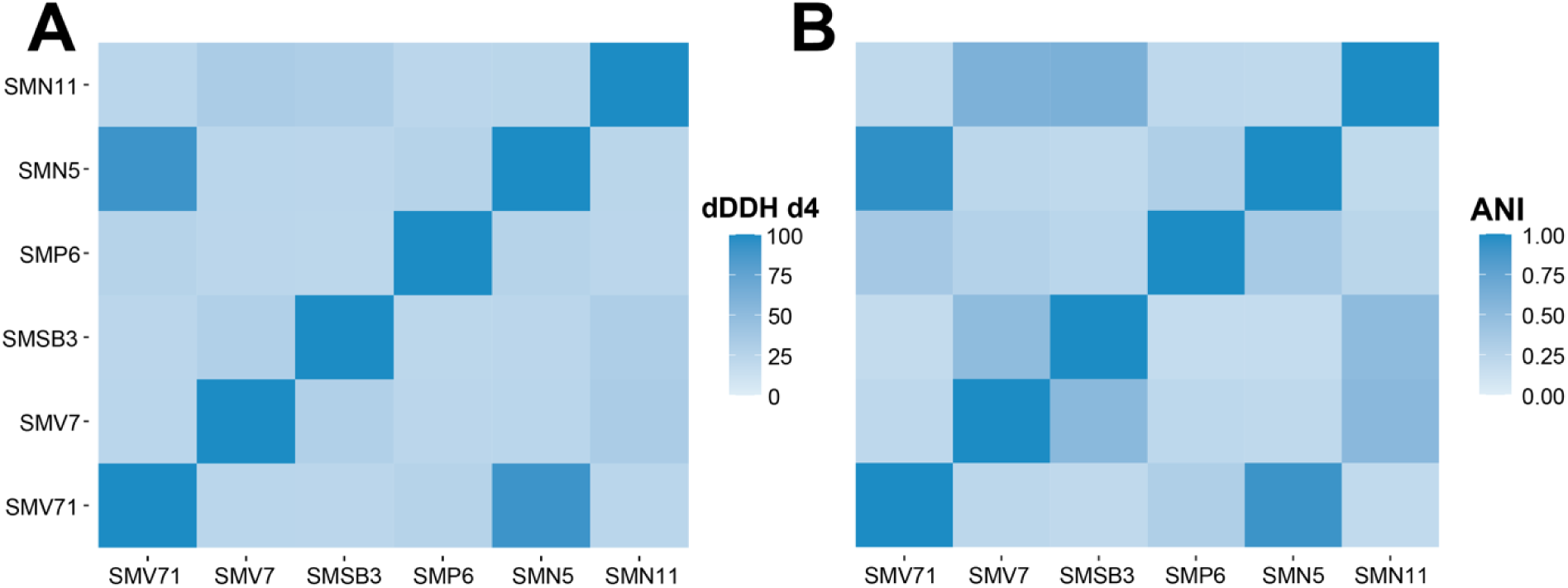
Whole genome comparison of the six *Pseudomonas* isolates. (A) digital DNA-DNA hybridization (dDDH) d_4_ values between strains. Average branch support was reported as 98.3% and *δ* value of 0.127. (B) Heat map of Average Nucleotide Identity (ANI) Hadamard analysis of strains using MUMmer-based ANI (ANIm).

**Table 2.**
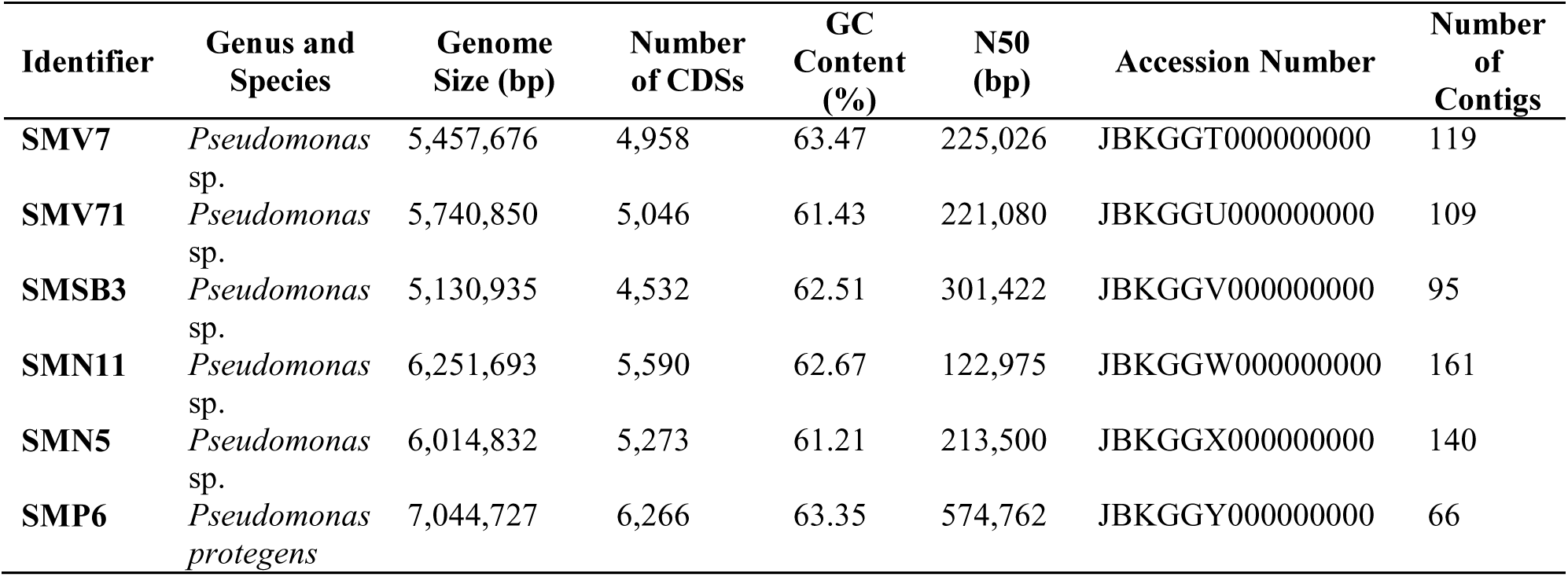
Statistics of the Whole-Genome Sequencing data for the six bacterial strains selected for the SynCom.

The other strains did not display digital DNA-DNA hybridization (dDDH) values with over 70% similarity to suggest that they had close enough phylogenetic relationships to match with *Pseudomonas* species already in the TYGS server (Morimoto et al. 2020). Strains SMN5 and SMV71 had a dDDH d_4_ value of 90.9% (Table 2), which suggested the strains are from the same species. Both dDDH d_2_ and d_6_ values corroborated this finding. To further assess the similarity of phylogenetic relationships between the strains, ANIm was used to compare the data in pairwise genomics. The same two strains, SMN5 and SMV71, displayed ANI similarity values of SMN5:SMV71 at 91.27% and SMV71:SMN5 at 95.62% (Fig. 1) and were of the same species determined by a threshold of 95%, considering genome completeness, size, and contiguity (Morimoto et al. 2020).

### Bacterial growth in the transwell system

OD_595_ measurements from the transwell system were used to compare the abundance of microbes in the synthetic community at the beginning versus the end of the experiment and to determine if the interactions with native microbial communities caused differences in growth rates, habits, or potential antagonistic interactions. The SynCom in the M9 minimal medium was used as a control group to analyze SynCom growth via OD_595_ readings. In contrast, SynCom the presence of the soil slurry was used to determine experimental effects. The overall growth of the SynCom in the control group displayed positive growth over time, while the treatment group had slight fluctuation before stabilizing (Fig. 2).

**Fig. 2.**
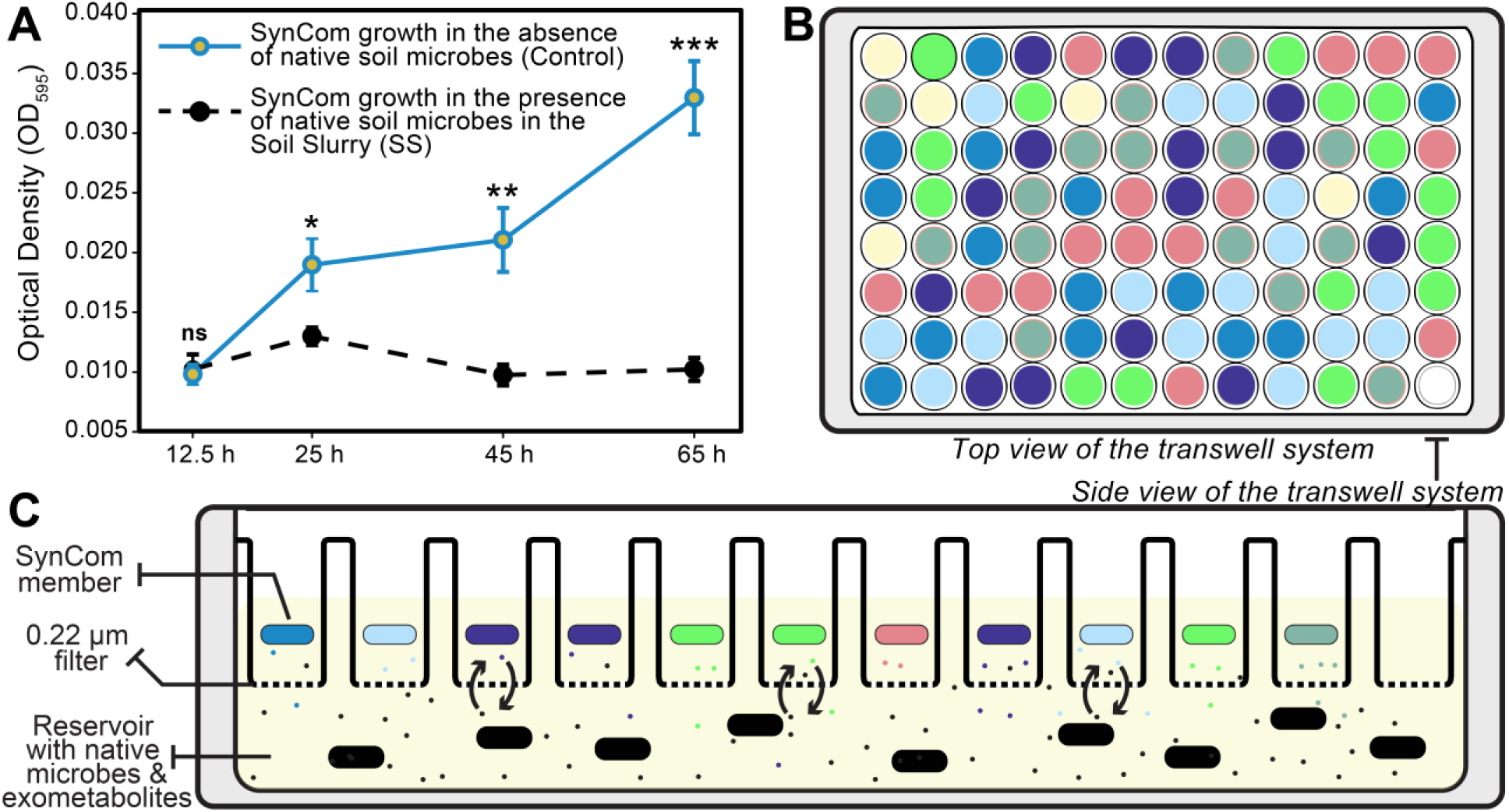
Overview of the transwell system for evaluating SynCom growth while exposed to native soil microbes. (A) SynCom growth (OD_595_) under control conditions and exposure to native soil microbes. The line on each bar represents ±SE. (Means of 2 experiments and 15 replicates (wells) per treatment). *, **, and *** represent significance by Student’s t-test at the 0.05, 0.01, and 0.001 probability level, respectively. Non-significant findings are represented by “ns.”. (B) The top view of the transwell system is where different colors represent the randomized well placement of six SynCom bacteria (red, yellow, green, light blue, dark blue, indigo) and wells with only M9 medium (yellow). (C) Side view of the transwell plate showing twelve wells containing SynCom bacteria constrained by a 0.22 µm filter. The SynCom bacteria and native soil microbes share media (yellow) and chemically interact but are spatially separated by the filter.

Over time, there were no considerable changes in the population sizes of the SynCom strains when interacting with the native microbiomes in the soil slurry (SS) (Fig. 3).

**Fig. 3.**
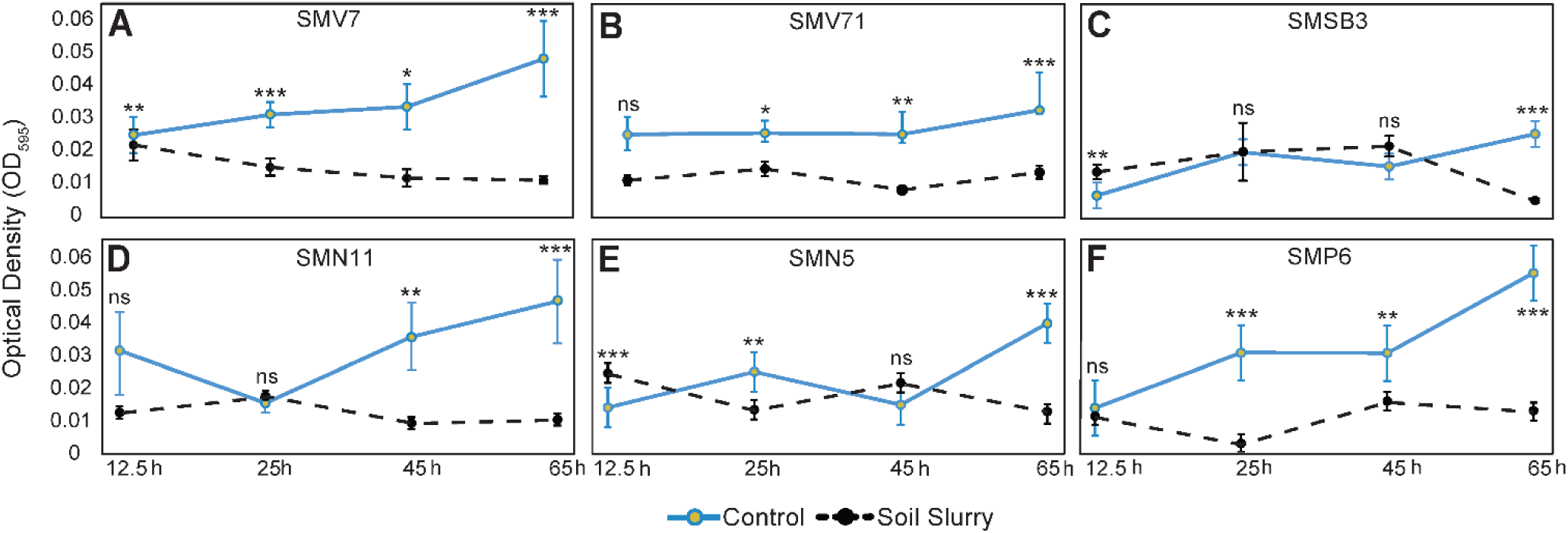
Bacterial growth assessment of synthetic microbial community (SynCom) members over 12.5, 25, 45, and 65 h. The blue/solid and black/dotted lines represent measurements of the control and soil slurry (SS) treatments, respectively. The graphs A-F shown represent the following *Pseudomonas* strains: A) SMV7, B) SMV71, C) SMSB3, D) SMN11, E) SMN5, and F) SMP6. The line on each bar represents ±SE. *, **, and *** represent significance by the Student’s t-test at the 0.05, 0.01, and 0.001 probability level, respectively. Non-significant findings are represented by “ns.”

The control and SS groups exhibited significant growth during the 12.5 to 25 h timepoint and a decline of the SynCom member of the SS treatment.

When assessed individually, the SynCom members showed different growth patterns among themselves and when compared to the SS treatment versus the control. For instance, strain SMV7 soil slurry treatment displayed an overall decrease in growth, while the control group of the strain showed progressive growth. SMN11 showed the largest decrease in growth from 12.5 h SS treatment compared to 65 h, with a 65.6% decrease. Strain SMV7 decreased by 54.7%, followed by SMV71 at 45.3%, SMSB3 at 20.7%, SMN5 at 7.53%, and finally SMP6 at 6.00%. Similarly, SynCom members in the control treatment showed an overall increase in growth, with some variations. On the other hand, the SS treatment measurements in between the strains remained relatively stable (Fig.3). When comparing the control to the treatment at 65 h, SMV7 had a 76.5% difference, SMV71 was 58.2%, 77.8% for SMSB3, SMN11 at 76.6%, SMN5 at 65.0%, and SMP6 with a 76.6% difference.

### Viability plating

Viability analysis used qualitative observation of strain morphological features and quantifying CFUs (colony-forming units) per mL. Throughout the four time points, viability results fluctuated from having significant to insignificant differences. At 65 h, differences in log-transformed values of CFUs mL^-1^ were not different enough to constitute a conclusion that viability between the control and treatment types was significant. Analysis of morphological features showed that for strain SMV71, colonies were visually observed to be smaller than when compared to the control plates. The control and treatment plates displayed similar or identical colony morphology over an additional 24-hour observation period. These findings support that bacteria survived or recovered despite the treatment showing significant declines in bacterial biomass, as suggested by the OD_595_ data.

Strains SMSB3, SMV7, SMP6, and SMN11 did not display differences in colony size between control and treatment plates. For trial 1, SMN5 showed persistent bacterial colony morphology but had no noticeable morphological differences in phenotype in subsequent trial observations. Differences in OD_595_ values but insignificant variation in viability suggest that bacterial cells persist in stress conditions and can produce similar numbers of CFUs mL^-1^, but at lowered biomass accumulation. Inferences may suggest that strain SMV71 was in a depressed state that, despite having similar numbers of CFUs mL^-1^, were slower growing and visually smaller than the bacteria from the same original strain solution in the control group.

### Flow cytometry analysis

Flow was conducted via comparison of unstained, live, and dead bacterial on two strains, one of which displayed persistent morphological characteristics during viability plating and one that did not (Fig. 4). The percentage of live cells for the control group of SynCom bacteria in the M9 minimal salts media for both SMP6 and SMV71 was 11.2% and 97.1%, respectively. The percentage of dead cells stained with PI, 1.8%, was measured to be lower in the control group than when in contact with the soil slurry for SMV71, 8.2%. SMP6 did not display statistically significant findings for the measures of live, dead, and unstained cell percentages.

**Fig. 4.**
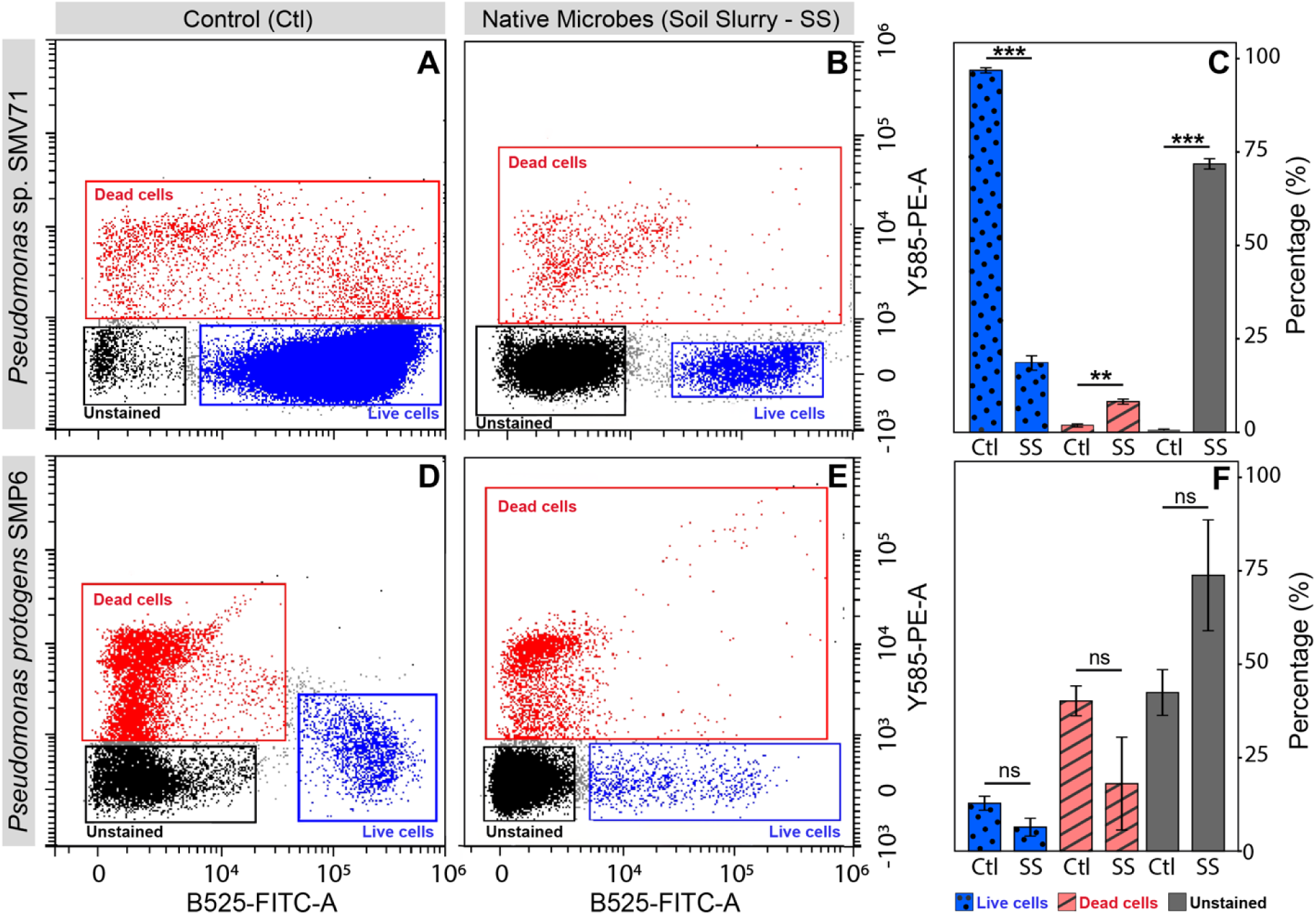
Flow cytometry data for *Pseudomonas* strains SMP6 and SMV71. Graphs A, B, D, and E display gates for bacterial cell groups: Live (blue), dead (red), and unstained (black) bacterial cell. (A) SMV71 cells for the control from the synthetic microbial community in M9 medium.(B) SMV71 cells from the SynCom in SS. (C) *Pseudomonas* sp. strain SMP6 as a control group from the SynCom in M9 medium. (D) Estimated cell percentages of for strain SMP6 in the native microbial slurry treatment. (F) Average cell type percentages for the control (Ctl) and native soil microbe (SS) groups for strain SMV71. Statistical analysis for Graphs C and F were performed using a paired t-test as specified in the methods. The line on each bar represents ±SE (Means of 2 experiments and 3 replicates). *, **, ***, or ns represent significant by Student’s t test at the 0.05, 0.01, and 0.001 probability level, and non-significance, respectively.

Checking stained live cells, we found that only 0.8% were unstained. Similarly, in the stained dead sample, we observed that only 1.1% of the total events were unstained. Under prime growth and nutrient conditions, these cultured bacteria samples displayed little to no inhibition in staining, which contrasted the results from the experimental control group in minimal medium and the treatment group.

### Carbon metabolism analysis

The Biolog EcoPlate was used to characterize the carbon metabolism of *Pseudomonas* spp. SMP6 and SMV71 over the course of a 72-hour time series. The measurements took place after 0 h and 65 h of exposure to native soil microbes in the transwell system. At 0 h native soil microbe exposure, SMP6 and SMV71 exhibited similar patterns of carbon utilization as evidenced by similar color change, measured as average well color development (AWCD), for the six compound classes (Fig. 5).

**Fig. 5.**
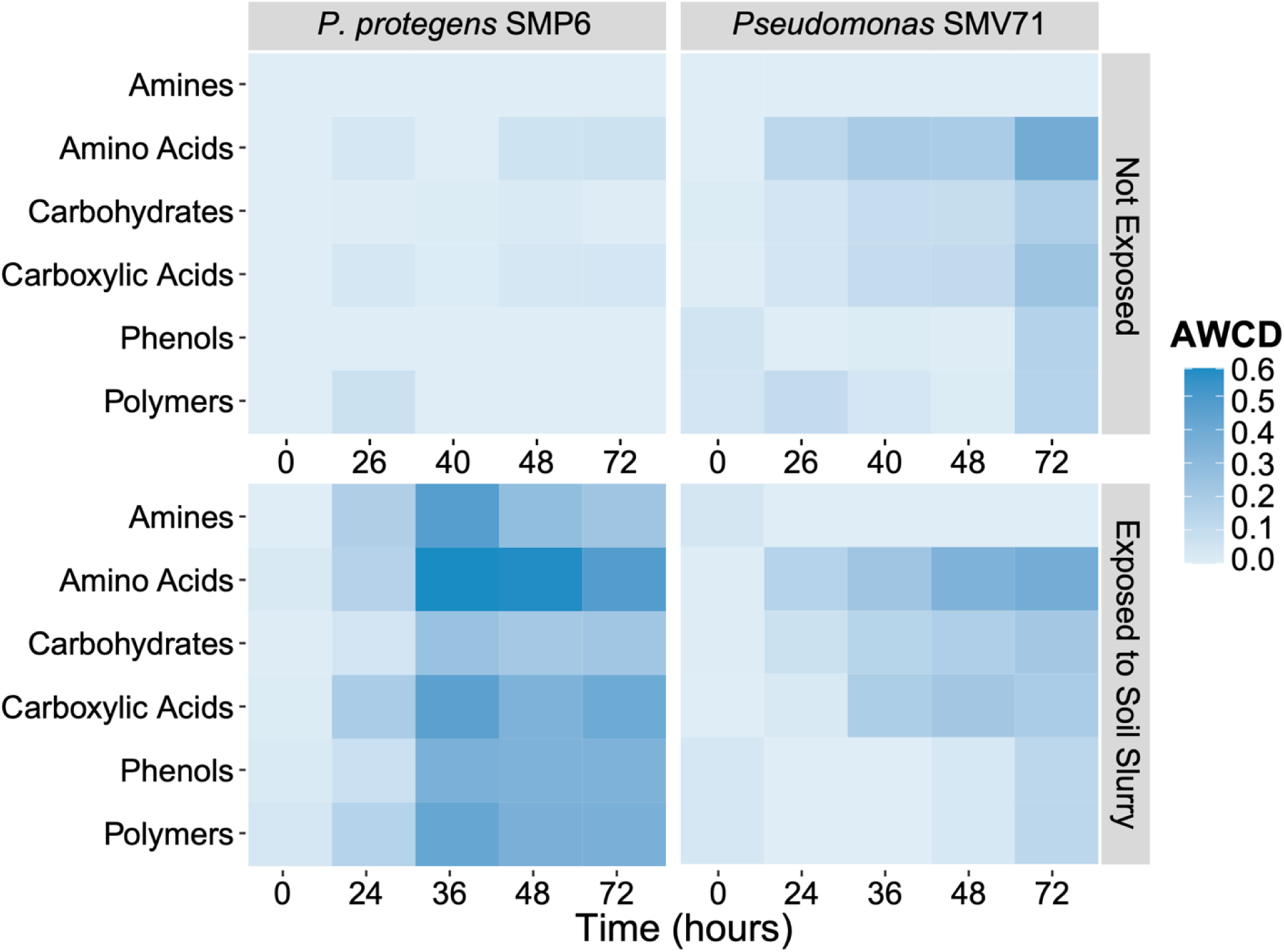
Analysis of bacterial carbon source utilization over a 72 h growth period with the Biolog EcoPlate^TM^. Carbon metabolism of *Pseudomonas protegens* SMP6 and *Pseudomonas* sp. SMV71 was measured as average well color development (AWCD) after 0 h and 65 h of exposure to native soil microbes. Means of three biological reps (plates) and three technical replicates (wells).

At 0 h, SMV71 exhibited slightly greater carbon compound metabolism from 40-72 h than SMP6, but overall carbon utilization was very similar between the bacterial species. The AWCD for all six compound classes for SMP6 was higher at 65 h than 0 h of soil slurry exposure, indicating that SMP6 responded through increased carbon utilization. In contrast, *Pseudomonas* SMV71 exhibited a consistent pattern and level of carbon utilization before and after soil slurry exposure.

Amino acids were the most utilized group of carbon for both SMP6 and SMV71. Of all the measured amino acids, L-asparagine was the most used at 0 h, followed by L-arginine and L-phenylalanine. After 65 h of soil slurry exposure, there was a greater diversity of amino acids utilized by both *Pseudomonas* spp., and L-asparagine remained the most used. At 0 h and 65 h, SMP6 and SMV71 also highly utilized the polymer Tween 40, D-galactonic acid γ-lactone (carbohydrate), and γ-aminobutyric acid (carboxylic acid). On the other hand, α-cyclodextrin, glycogen, D-cellobiose, β-methyl-D-glucosidase, α-D-lactose, α-ketobutyric acid, and most amines and phenols were poorly utilized. The 65 h exposure to the native soil microbes also altered bacterial utilization of several carbon compounds. Before soil exposure, SMP6 and SMV71 primarily utilized the carbohydrate D-xylose, whereas, after 65 h soil exposure, D-mannitol became one of the most highly utilized carbohydrates. Additionally, the utilization of γ-aminobutyric acid drastically increased for SMP6 and SMV71 after exposure to the soil slurry. SMP6 and SMV71 also exhibited differences in carbon metabolism. At both time points, the growth of SMP6 was favored in the presence of the polymer Tween 80, whereas SMV71 did not utilize this compound. Conversely, D-galacturonic acid (carboxylic acid) was an important carbon source for SMV71 but was not highly utilized by SMP6.

### Secondary metabolite prediction

Whole-genome sequences were analyzed for the prediction of secondary metabolites using biosynthetic gene clusters (BGCs) via antiSMASH version 7.1.0. This analysis provides insights into the genetic potential of each strain to produce secondary metabolites but does not indicate actual metabolic activity. Experimental confirmation through transcriptomics (e.g., RNA sequencing) or metabolomics would be required to determine whether these biosynthetic pathways are actively expressed under specific conditions.

The biosynthetic potential varied across the six SynCom strains (Fig. 6). *Pseudomonas protegens* SMP6 displayed the highest number of predicted BGCs, totaling 27, and exhibited the most diverse range of BGC classes compared to other strains. SMN5 and SMV71 followed with 22 and 20 BGCs, respectively, while strain SMSB3 had the fewest detected BGCs at 12.

**Fig. 6.**
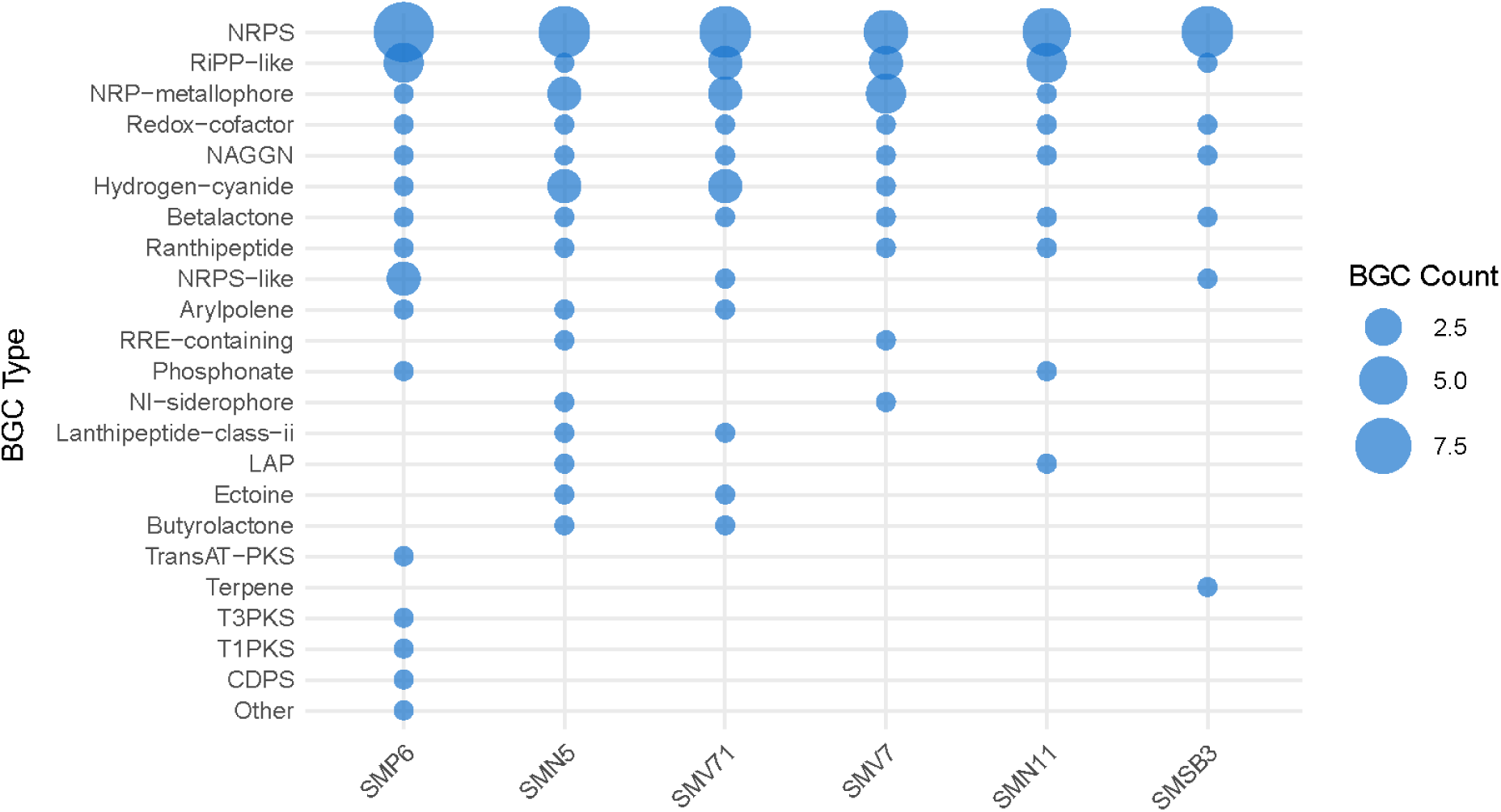
Biosynthetic Gene Clusters (BGCs) predicted in each bacterial strain used in the synthetic microbial community were identified using antiSMASH software. Strains were arranged from left to right based on the total number of predicted BGCs, listed in descending order from *Pseudomonas protegens* strain SMP6 (highest) to *Pseudomonas* sp. strain SMSB3 (lowest). The quantification of BGCs is based on the number of distinct BGC types identified through bacterial genome analysis for each strain.

Nonribosomal peptides (NRPS) were the most abundantly identified BGC class across all six strains. RiPP-like and NRPS-metallophore BGCs were also conserved throughout the strains, though to a lesser extent. Additionally, each strain contained a single BGC associated with the biosynthesis of β-lactone, NAGGN, and redox-cofactor compounds.

AntiSMASH also predicted specific clusters with varying confidence levels. Strain SMV71 had no predicted clusters with 100% homology to known BGCs. However, it was predicted to encode a BGC for hydrogen cyanide production, as well as the nonribosomal peptide kolossin. Strain SMSB3 was predicted to harbor a terpene BGC associated with carotenoid biosynthesis. Miscellaneous BGCs were identified in SMN11 and SMN5. Additionally, hydrogen cyanide biosynthetic clusters were predicted in strains SMV71, SMN5,and SMP6. Notably, antiSMASH predicted a hybrid T1PKS/RiPP-like BGC encoding an NRPS-metallophore and a Type III polyketide synthase (T3PKS) cluster predicted to synthesize 2,4-diacetylphloroglucinol.

These findings illustrate the biosynthetic potential of each strain based on genome annotation, but further studies are necessary to determine whether these gene clusters are actively expressed and producing secondary metabolites under relevant environmental conditions.

### Shotgun metagenomic analysis for soil slurry microbiome

The complete taxonomic diversity of both control and soil slurry-treated samples, as determined through metagenomic profiling based on relative abundance values, were provided in the supplementary Table 1. The twenty most abundant species identified in the soil slurry at 65 h were visualized using the heatmap (Fig. 7A). Hierarchical clustering reveals distinct microbial composition shifts over time, with notable differences in species abundance patterns. The community exposed to the soil slurry was dominated by species such as *Comamonas thiooxydans*, *Delftia acidovorans,* and a few *Pseudoxanthomonas* sp. sustained presence and potential enrichment of the *Pseudomonas urmiensis*, P*seudomonas putida, Pseudomonas umsongensis* and *Pseudomonas* sp. *s*train C1C7 at 65 h, suggested its persistence within the microbial community.

**Fig. 7.**
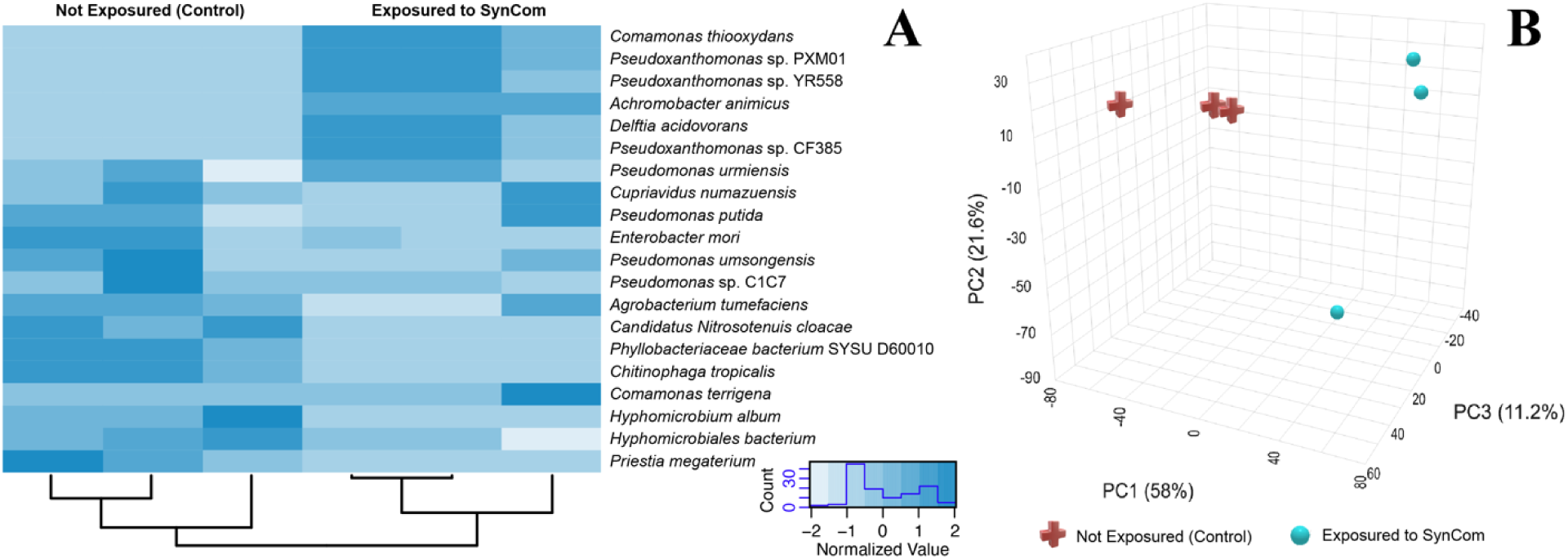
Taxonomic abundance and pathway composition of soil slurry microbiome at 0 h and 65 h. A) Heatmap of the 20 most abundant microbial species identified by MetaPhlAn 4.0. The color gradient represents species abundance, from higher relative abundance to lower abundance. Differences in species composition between the initial timepoint and after exposure to soil slurry highlight shifts in the microbial community over time. B) 3D Principal Component Analysis (PCA) plot of pathway abundance profiles in the soil slurry microbiome at 0 h (red) and 65 h (blue). Pathway abundance was determined using HUMAnN v3.0 and visualized in MicrobiomeAnalyst. The three principal components, PC1 (58%), PC2 (27.6%), and PC3 (11.2%), capture most of the variation in pathway composition.

To assess functional potential changes, we analyzed pathway abundance using HUMAnN v3.0. A total of 10951 pathways were identified across exposed and non-exposed soil slurry. (Supplementary Table 2). Top five pathways involving fatty acid metabolism (PWY-5136: fatty acid and beta;-oxidation II plant peroxisome, PWY-7663: gondoate biosynthesis (anaerobic), amino acid biosynthesis (VALSYN-PWY: L-valine biosynthesis), and stress response related metabolic pathways including PWY0-1586: peptidoglycan maturation (meso-diaminopimelate) and PWY-5973: cis-vaccenate biosynthesis were identified. After exposure to the soil slurry, there was an apparent increase in the abundance of pathways associated with fatty acid metabolism and cell wall remodeling, possibly in response to increased stress. In PCA, PC1 (58%), PC2 (21.6%), and PC3 (11.2%) accounted for the majority of the variance in pathway abundance (Fig. 7B). The distinct clustering of the 0 h and 65 h samples suggested significant shifts in microbial metabolic functions over time, indicating changes in the community’s functional potential following exposure to the soil slurry.

## Discussion

Developing synthetic microbial communities holds promise in inducing host immunity and defense responses. Understanding the interactions between microbes and their appointed environments and determining compatible microbes that provide specialized functions is key for ensuring SynCom stability, longevity, and success (Martins et al. 2023). This study found that a SynCom that was composed of six *Pseudomonas* spp. demonstrated compatibility, with some members displaying persistent traits. Compatibility between SynCom microbes is vital for long-term applications because microbial instability impairs successful SynCom implementation in an applied environment (Shayanthan et al. 2022). Based on persistent characteristics, some of the SynCom members investigated in this study are promising candidates for future applications *in situ*.

The transwell system is specialized for interpreting metabolic and chemical interactions between SynCom microbes and reservoir without physical interaction (Chodkowski and Shade, 2024). Analyzing microbial interactions is valuable to improving the longevity and stability of a SynCom. Additionally, this analysis unveils bacterial persistence as a response to environmental or antagonistic stresses. For instance, native rhizosphere microbes may recognize introduced microbes as antagonistic, potentially reducing the effectiveness of lab findings when applied in field conditions (Thomas and Sekhar, 2016). In individual OD_595_ tests, the SS groups exhibited stable or reduced growth fluctuations when in contact with native soil microbes, when compared to the initial growth at 0 h, when the test was set up. Throughout different growth phases, *Pseudomonas* spp. undergo diverse physiological changes in response to interactions with other cells and environmental stresses. Studies suggest that persister cells are most common in the stationary-phase *Pseudomonas* spp., where sessile, or biofilm-associated bacteria, predominate (Spoering and Lewis, 2001). Bacteria that grow in the presence of antagonism are generalized as resistant (Wood et al. 2013). The SynCom bacteria do not demonstrate this characteristic, further supporting the fact that the bacteria are persistent rather than resistant.

For strains exhibiting a reduction in OD_595_, the decline in biomass accumulation suggests antagonism or competition between the SynCom and native bacteria for resources (Sun et al. 2023). Despite lowered biomass in the SS treatment, similarities in CFU counts show that the bacteria are viable but not growing the same as the control group. These observations may offer insights into SynCom stability in applied environments, where stress tolerance mechanisms and persistence traits are critical for long-term bacterial survival and functionality.

Specialized bacterial defense mechanisms affect the population dynamics by facilitating resource competition and antagonism between the SynCom and native soil microbes. *Pseudomonas* species are known to utilize antibacterial toxins via specialized secretion systems to combat antagonism (Peterson et al. 2021). Secondary metabolites are common amongst soil bacteria and may lead to competition between surrounding organisms that overlap in the same niche (Peterson et al. 2021). Bacterial competition for resources against adversary bacteria is augmented by nutrient-limited environments (Hibbing et al. 2010). Persistence in bacteria is suggested to be governed by environmental conditions and growth phase, particularly in stationary growth phases where environmentally limiting factors such as nutrient availability are significant (Kunnath et al. 2024). Bacteria compete for the minimally supplied resources in a transwell system, utilizing competitive dynamics such as inhibitory compounds.

In the face of stress and exposure to antagonism, bacterial persister cells can enter a dormant state of reduced or halted growth and metabolism to protect themselves against intoxication through specific physiological changes (Harms et al. 2016). Not all bacteria within the population show persistent traits, while the majority are still susceptible to the stressor. Depending on the circumstance and growth stage, each SynCom strain can have persister cells. Frequently utilized protective adaptations include altered gene expression or physiological patterns, including altered barrier membrane function and biofilm formation (Shree et al. 2023). Bacterial cells that defend themselves through this dormancy exhibit decreased growth before returning to their normal physiological state (Harms et al. 2016). SMV71 colonies initially displayed similar numbers of CFUs mL^-1^, but with smaller bacterial colonies in the soil microbial slurry treatment group compared to the control. After being introduced to high-nutrient media, the colonies recovered from a persistent state. Recovery is an important part of bacterial persistence, as limited growth and inhibited metabolic processes are not enough to enable bacterial survival under stress conditions (Pan et al. 2023).

Flow cytometric analysis can assess SynCom viability after exposure to native soil microbes. Although having a comparable proportion of stained cells is cardinal, disparities between staining among groups can provide insight into bacterial persistence. For both SMP6 and SMV71, the treatment exposed to the soil slurry inhibited bacterial staining (a difference of over 71% in SMV71 and over 31% in SMP6), which was not observed in the control group. Biofilm formation, a complex polymetric structure, is a protective physical barrier against environmental stresses, predation, and chemical attacks by both biotic and abiotic competition (Shree et al. 2023). Biofilms can prevent staining by serving as a barrier of bacterial “protection” against extracellular influences, thus inhibiting visualization of cell viability. Therefore, greater amounts of unstained cells in flow cytometric analysis were detected in both Pseudomonas spp. following soil slurry exposure supports that biofilm formation may be involved as a survival tactic in SMP6 and SMV71.

Overall, the carbon utilization traits observed through the Biolog EcoPlate analysis demonstrate persistence in the tested strains. SMV71 maintained a consistent carbon utilization at 0 h and 65 h of soil slurry exposure, while SMP6 responded through increased carbon utilization. Despite these differences, both strains responded through resource generalism and utilized a more diverse array of amino acid and carbohydrate sources after exposure to the native soil microbes. Bacterial dormancy is not always associated with a decreased metabolism but rather a diversification of nutrient sources and active metabolic pathways (Greening et al. 2019). Even during low-growth and quiescent states, microbes often accumulate carbon stores in preparation for improved conditions and exiting dormancy (Rittershaus et al. 2013). Protein turnover may also increase in dormant cells, especially while exiting dormancy, as exhibited in SMV71 and SMP6 through increased utilization of amino acids after 65 h soil slurry exposure (Rittershaus et al. 2013). Diversification of carbon source utilization following exposure to the native soil microbes, observed in SMV71 and SMP6, is a persistence trait.

Analyzing the temporal changes in carbon source utilization can also provide insight into the response of SMP6 and SMV71 to the native soil microbiome. For example, the utilization of γ-aminobutyric acid (GABA) in both bacteria significantly increased after 65 h of soil slurry exposure. The GABA shunt is a metabolic pathway that indirectly converts GABA into succinate, fueling the tricarboxylic acid cycle, raising cellular pH, and generating reduced power. This pathway is present in many bacteria, including some *Pseudomonas* spp., and has been implicated as an important facet in the bacterial response to acid, osmotic, anaerobic, and oxidative stresses (Feehilly and Karatzas 2012). Although this pathway was detected in low abundance in the soil slurry, the EcoPlate analysis supports that the GABA shunt may be an important pathway in the response of some SynCom microbes to the native soil microbes.

Biosynthetic gene clusters serve many functions in bacterial survival and persistence, supporting competitive advantages (Dilshad et al. 2024). Secretion of inhibitory or antibiotic compounds is a pivotal characteristic of SynCom members. Ribosomally-synthesized and post-translationally modified peptides (RiPP and RiPP-like compounds) serve as precursor peptides for immunity determinants and were highly conserved across all of the SynCom strains (Peterson et al. 2021). NRPs and polyketides (PKs) synthesize antimicrobial compounds and provide self-protection from their bioactive secretions (Peterson et al. 2021). The NRP *Pseudomonas fluorescens* Pf-5 pyoverdine is an iron siderophore capable of starving competition of iron. This makes Pf-5 pyoverdine an important component of antimicrobial compounds and a stabilizer for the biofilm polysaccharide matrix (Hartney et al. 2013; Khasheii et al. 2021).

Each strain was predicted to synthesize β-lactone, NAGGN, and redox-cofactor. β-lactones are often enzyme inhibitors, including effective antimicrobials (Schaffer et al. 2017). N-acetylglutaminylglutamine amide, or NAGGN, acts in response to osmotic stress (D’Souza-Ault et al., 1993). Redox cofactor is present in each strain and promotes energy acquisition, suggesting metabolic promise in promoting respiration-induced biofilm formation (Matrín-Rodrríguez, 2023; Mai-Prochnow et al. 2015).

The synthesized compounds predicted from the various BGCs display valuable SynCom composition and persistence functions. For instance, hydrogen cyanide is a cyclic lipopeptide, which is a common antimicrobial metabolite present across a large spectrum of *Pseudomonas* species and has been shown to have inhibitive properties against fungi, oomycetes, bacteria, viruses, and protozoans (Masschelein et al. 2017; Zhao et al. 2019). Additionally, these cyclic lipopeptides act as biosurfactants and affect biofilm formation and motility (Geudens and Martins, 2016). 2,4-diacetylphloroglucinol, found in SMP6, can disrupt bacterial cell membranes and ultimately reduce cell viability (Bangera et al. 1999).

These different secondary metabolites give each strain unique qualities that aid in bacterial defense and offense, in addition to functions in host defense and offense. These traits support competitive fitness and microbial persistence under stress conditions. BGCs that synthesize compounds with antimicrobial properties and support biofilm formation give hosts a competitive advantage.

Metagenomic diversity analysis revealed significant shifts in microbial community composition following soil slurry exposure. Notably, the enrichment of *Pseudoxanthomonas* significantly increased after exposure, suggesting the genus’s versatility in adapting to changing environmental conditions (Choi et al. 2013). Additionally, the persistence of particular *Pseudomonas* species highlights their potential to enhance the longevity of the SynCom under fluctuating environmental conditions. The distinct clustering at two different time points indicates temporal shifts in microbial metabolic functions, suggesting changes in community functional potential in response to soil slurry exposure. These shifts correspond with increased abundance in pathways related to fatty acid metabolism and cell wall remodeling, which may indicate stress-induced metabolic adaptations. Functional Pathway analysis suggests that SynCom bacteria utilized fatty acid degradation pathways to adapt to environmental stress. The EcoPlate analysis supports this finding because following soil slurry exposure, fatty acid-containing Tween 40 and 80 increased for SMP6, and Tween 40 utilization also increased for SMV71.

Additionally, within the native soil slurry, GLUDEG-I-PWY: GABA shunt was detected in both the 0 h and 65 h samples but showed low abundance. The low representation of this pathway in both groups suggests that GABA metabolism may not be a primary survival strategy under the treatment conditions, despite its known role in stress response and metabolic flexibility. Cell wall remodeling is another crucial survival strategy for bacteria, particularly in stressful environments (Mueller and Levin, 2020). The upregulation of pathways associated PWY0-1586: peptidoglycan maturation and PWY-5973: cis-Vaccenate Biosynthesis may reflect a microbial strategy to enhance structural integrity or modulate interactions with other microbes and environmental stressors. These changes were due to the exchange of microbial metabolites, given the fact that transwell system does not allow for physical contact.

### Conclusion

Retaining microbiota that can persist in their implemented environment is vital to improving the stability and longevity of SynComs. Diversifying species in SynCom design can incorporate higher variability in bioactive metabolites but also has a greater number of interactions with interfering microorganisms, pathogens, and members of the SynCom itself (Martins et al. 2023). Ensuring SynCom compatibility and stability in recipient soils is crucial for effective biocontrol and plant immune defense while preventing dysbiosis. Supporting SynCom bacteria in recovering from dormant or persistent states enables them to fulfill their designed roles, such as enhancing host defense or promoting growth.

## Funding information / Acknowledgements

This work was partially supported by the USDA National Institute of Food and Agriculture, Hatch project 1024881, NIFA grant project N° 2022-51300-43051. The authors also acknowledge the University of Florida Interdisciplinary Center for Biotechnology Research (ICBR)/ Cytometry Core (RRID:SCR_019119) for the assistance with the flow cytometry assay.

